# Formation of amyloid fibrils by the regulatory 14-3-3ζ protein

**DOI:** 10.1101/2023.05.31.543065

**Authors:** Darius Šulskis, Mantas Žiaunys, Andrius Sakalauskas, Vytautas Smirnovas

## Abstract

The 14-3-3 is a highly conserved adaptor protein family with multi-layer functions, abundantly expressed in the brain. The 14-3-3 proteins modulate phosphorylation, regulate enzymatic activity and can act as chaperones. Most importantly, they play an important role in various neurodegenerative disorders due to their vast interaction partners. Particularly, the 14-3-3ξ isoform is known to co-localize in aggregation tangles in both Alzheimer’s and Parkinson’s diseases as a result of protein-protein interactions. These abnormal clumps consist of amyloid fibrils – insoluble aggregates, mainly formed by amyloid-β, tau and α-synuclein proteins. However, the molecular basis of if and how 14-3-3ξ can aggregate into amyloid fibrils is unknown. In this study, we describe the formation of amyloid fibrils by 14-3-3ξ utilizing a comprehensive approach that combines bioinformatic tools, amyloid-specific dye binding, secondary structure analysis and atomic force microscopy. The results presented herein characterize the amyloidogenic properties of 14-3-3ξ and imply that the well-folded protein undergoes aggregation to β-sheet-rich amyloid fibrils.

## Introduction

14-3-3 are acidic proteins that first were discovered in bovine brain extracts and titled after their migration position in two-dimensional electrophoresis^1^. Afterwards, these proteins were identified as a highly conserved protein family that is found in all eukaryotes^2^. They have been observed to interact with various kinases and enzymes via recognizing phosphorylation sites^3^. Through the vast interaction network, 14-3-3 proteins regulate the activity and stability of proteins, control localization, and facilitate protein-protein interactions (PPI)^4^.

There are seven known mammalian isoforms (β, γ, ε, η, ζ, σ and τ/θ) of 14-3-3 proteins^5,6^, all of which are dimeric. Each monomer is composed of nine antiparallel α-helices and a disordered C-terminal tail, that is suggested to have an autoinhibitory role against non-specific interactions^7^. Their overall structure resembles a clamp shape with a conserved amphipathic grove, which is used as an accessible binding site for a plethora of proteins^8^. The different interaction specificity of isoforms comes from individual domain movements that ensure flexible adaptations of the binding surfaces^9^.

14-3-3 proteins are found in various human tissues^10^, despite being expressed most abundantly in the brain tissues^11^. Most prominently, they regulate important physiological functions such as cell survival, differentiation, migration, apoptosis and ion channel regulation^12^. Due to their involvement in numerous crucial roles in the nervous system, they are also associated with neurodegeneration disorders^13^. Different 14-3-3 isoforms have been found in amyloid plaques (neurofibrillary tangles^14^ and Lewy bodies^15^), which are indications of ongoing neurodegenerative diseases^16,17^. Although, they seem to have conflicting roles, as one report demonstrates their ability to facilitate the formation of microtubule-associated Tau protein fibrils^18^, while another study shows their potential to disrupt Tau liquid-liquid phase separation and thus inhibit amyloid aggregation^19^. The increased chaperone-like activity of 14-3-3 proteins was observed with Parkinson’s disease-related α-synuclein protein^20^. 14-3-3 inhibited α-synuclein aggregation *in vivo*^21^ and rerouted its aggregation pathway through binding to intermediate α-synuclein species^22^. Due to these important associations with amyloidogenic and other protein partners, the 14-3-3 family is considered to have a novel therapeutic potential for neurological diseases^16^.

While 14-3-3 proteins play a prominent role in neurodegenerative diseases because of their synergy with various partners, it has never been investigated whether they may themselves be susceptible to amyloid aggregation. Previously, there have been several indications that 14-3-3 might be amyloidogenic proteins, since it is known that 14-3-3 share physical and functional homology with α-synuclein^23^, particularly in relation to its non-amyloid-β component and C-terminal tail amino-acid sequence homology^24^. In addition, it has recently been shown that the concentration of 14-3-3ζ isoform in cerebrospinal fluid increases during the early stages of Alzheimer’s disease^25^, indicating a potential link with the disorder pathogenesis.

In this study, we investigated whether the 14-3-3ζ isoform can aggregate into amyloid fibrils. The preliminary computational data showed that the 14-3-3ζ amino acid sequence contains six potential aggregation-prone sites. Subsequently, we monitored protein aggregation using various amyloid-specific dyes and observed gradual, amyloid aggregation over seven days. The secondary structure analysis confirmed that the exclusively helical protein underwent structural changes and formed β-sheets, akin to the ones found in amyloid fibrils. Finally, AFM imaging confirmed that 14-3-3ζ aggregates consisted of short, curvy, fibril-like structures.

## Experimental Procedures

### Protein expression and purification

The 6xHis-SUMO-14-3-3ζ construct used in this work (kind gift of prof. B.M. Burmann) was derived from GST-14-3-3ζ (Addgene #13278)^26^. The plasmid containing 6xHis-SUMO-14-3-3ζ was chemically transformed into One Shot™ *E. coli* BL21 Star™ (DE3) (Fisher Scientific) cells. The transformed cells were grown at 37°C in LB medium containing kanamycin (50 mg/ml) until an optical density at 600 nm ≈ 0.7 was reached. Expression was induced by the addition of 0.4 mM isopropyl-β-d-thiogalactopyranoside (Fisher Scientific), and the cells were left to grow overnight at 25°C. Cells were harvested by centrifugation at 6000 x g for 20 min at 4°C and subsequently resuspended in 50 ml of lysis buffer (25 mM Hepes/NaOH, 1 M NaCl, and 10 mM imidazole (pH 7.5)). The suspension was lysed with a Sonopuls (Bandelin) homogeniser (10 s on, 30 s off, 30% power, total time 30 min). Cell debris was removed by centrifugation at 18,000 x g for 45 min at 4°C, and the supernatant was applied to a Ni^2+^ Sepharose 6 Fast Flow (GE Healthcare)–loaded gravity column, followed by stepwise elution with 20 ml of lysis buffer supplemented with 100 and 500 mM imidazole, respectively. Fractions containing the 6xHis-SUMO-14-3-3ζ protein were dialyzed against phosphate-buffered saline (PBS, pH 7.4), and the 6xHis-SUMO tag was removed by enzymatic cleavage using human Sentrin-specific protease 1 (SENP1) catalytic domain (derived from pET28a-HsSENP1, that was a gift from Jorge Eduardo Azevedo (Addgene plasmid #71465) at 4°C overnight^27,28^. The cleaved proteins were applied again to a Ni^2+^ column, and the flow-through was collected. The proteins were concentrated using Amicon centrifugal filters [10k Molecular weight cut-off (MWCO), Merck Millipore] and purified further by size exclusion chromatography (Superdex 75, GE Healthcare) in PBS.

### Aggregation-prone sequence analysis

Predicted 14-3-3ζ aggregation-prone regions were calculated using three different predictors: PASTA 2.0^29^, FoldAmyloid^30^ and AGGRESCAN^31^. The default parameters were used for each prediction. Briefly, in PASTA 2.0 90% sensitivity and -2.8 energy cutoff were used. In FoldAmyloid, aggregation sites were positive, if five successive amino acids with a score above 21.4 were detected. AGGRESCAN identified hot spots of aggregation, whenever the window of five amino acids had an amino-acid aggregation-propensity value higher than -0.02. The raw data of predictors results are presented in the supplement (Fig. S1).

### 14-3-3ζ aggregation

The purified 14-3-3ζ was diluted to 100 μM using a PBS pH 7.4 solution and distributed to 2.0 mL non-binding test tubes (400 μL volume, each test tube contained two 3 mm glass beads). The samples were then incubated at 37°C with constant orbital 600 RPM agitation. Every 24 hours, three samples were taken for analysis.

### Fluorescence and absorbance assays

For dye-binding assays, thioflavin-T (ThT), 8-anilinonaphthalene-1-sulfonic acid (ANS) and Congo red (CR) powders were dissolved in PBS buffer solutions and filtered using 0.22 μm syringe filters. The final concentrations of the dye solutions were set to 20 μM (ThT – ε_412_ = 23 250 M^-1^cm^-1^, ANS – ε _351_ = 5 900 M^-1^cm^-1^, CR – ε_486_ = 33 300 M^-1^cm^-1^) based on their specific absorbance spectra, which were scanned using a Shimadzu UV-1800 spectrophotometer. Amytracker630 stock solution was diluted 40 times using PBS buffer prior to use. The prepared dye solutions were stored at 4°C in the dark.

The dye solutions were combined with either PBS, the initial 14-3-3ζ solution or the incubated 14-3-3 ζ solution in a 1:1 ratio. The mixtures were then incubated for 10 minutes in the dark. The fluorescence spectra of ThT, ANS and Amytracker630 were scanned using a Varian Cary Eclipse spectrofluorometer with 10 nm excitation and emission slits, 1s averaging time and 1 nm intervals (ThT – 440 nm excitation and 460 – 540 emission range, ANS – 370 nm excitation and 420 – 560 emission range, Amytracker630 – 480 nm excitation and 580 – 680 nm emission range). The absorbance of CR was scanned from 200 nm to 800 nm using a Shimadzu UV-1800 spectrophotometer. All spectra were corrected using control samples, which did not contain the dye molecules.

### Light scattering assay

Sample right-angle light scattering was scanned with a Varian Cary Eclipse spectrofluorometer, using 600 nm excitation and emission wavelengths with 2.5 nm slit widths, 1s averaging time.

### FTIR experiments

The aggregated 14-3-3ζ samples (2 mL volume, 100 μM initial protein concentration) were centrifuged for 15 min at 12 000 RPM. Afterwards, the supernatant was carefully removed and replaced with 500 μL D_2_O with 400 mM NaCl (the addition of NaCl improves aggregate sedimentation). The centrifugation and resuspension procedure was repeated 3 times. After the final centrifugation, the aggregate pellet was resuspended into 50 μL of D_2_O with NaCl. The suspension was then scanned as described previously^32^ using a Bruker Invenio S FTIR spectrometer. Data analysis was carried out with GRAMS software. D_2_O and water vapor spectra were subtracted from the sample spectrum, which was then normalized between 1700 cm^-1^ and 1600 cm^-1^.

To acquire the FTIR spectra of monomeric 14-3-3ζ, the buffer solution (PBS, pH 7.4) was exchanged into D_2_O with 400 mM NaCl using a 10 kDa protein concentrator. The protein solution was diluted to 100 μM using the D_2_O solution, placed in the concentrator (400 μL volume) and centrifuged for 10 minutes at 9000 RPM. The concentrated protein solution (∼50 μL) was then diluted to the original volume with the addition of 350 μL D_2_O. This centrifugation and dilution procedure was repeated 4 times. The final concentrate was diluted to 100 μL and used for FTIR analysis. The spectra were obtained and analyzed the same as the 14-3-3ζ aggregate sample.

### Circular dichroism spectroscopy

All measurements were performed on a Jasco J-815 spectrometer at room temperature. Briefly, either a freshly prepared monomeric solution of 100 μM 14-3-3ζ (PBS, pH 7.4) or aggregated sample was transferred to a 0.1 mm quartz cuvette. Spectra were measured at 1 nm data pitch from 190 to 250 nm with a bandwidth of 2 nm and scanning speed of 50 nm/min. The final spectra of each sample were averaged from 3 scans with the buffer background subtracted. Analysis and visualization of spectra were done in Spectragryph software (http://spectragryph.com).

### AFM measurements

Atomic force microscopy imaging was done similarly as previously described^33^. In brief, 40 μL sample of aggregated 14-3-3ζ was placed on freshly cleaved mica that was functionalized using 40 μL of APTES ((3-Aminopropyl)triethoxysilane), incubated for 5 minutes. Then the mica was washed with 2 mL MiliQ water and dried gently under a stream of air. High resolution images (1024 × 1024) were collected using Dimension Icon (Brucker) AFM operating in tapping mode (Tap300AI-G silicon cantilever (40 N^-1^m^-1^, Budget Sensors). AFM image flattening and fibril analysis were done using Gwyddion v2.57^34^.

## Results and Discussion

### 14-3-3ζ contains aggregation-prone regions

14-3-3ζ have been located in various plaques of aggregated proteins in different neurodegenerative diseases. It is possible that strong protein-protein interactions, lead to their co-localization or even co-aggregation with amyloidogenic proteins (Fig. 1A). As 14-3-3ζ and other family members share similarities to α-synuclein^23,24^, which is a main aggregated component of Lewy bodies^20^, they can be suspected to be amyloid-forming proteins. Hence, we analyzed the 14-3-3ζ amino acid sequence to identify whether 14-3-3ζ contains aggregation-prone regions. We used three different predictors of aggregation-prone sites: FoldAmyloid^30^, AGGRESCAN^31^ and PASTA 2.0^29^. The aim was to identify whether 14-3-3ζ has potential aggregation regions and not to quantify them, therefore each positive site from at least two predictors was characterised as a hit. Overall, results from AGGRESCAN and FoldAmyloid indicated consistent aggregation-prone regions in six sites, whereas PASTA 2.0 only showed three (Fig. 1B). We identified that the six sites, which overlap between different algorithms, are found in five α-helices of 14-3-3ζ (Fig. 1 B,C). These results matched with previously predicted 14-3-3ζ aggregation regions in α2, α3 and α8 helices by PASTA 2.0 and AMYLPRED2^24^, while also suggesting potential involvement of α4 and α6. Since different predictors indicated that 14-3-3σ might aggregate, in order to verify that, we examined 14-3-3ζ aggregation properties with various biophysical methods.

**Figure 1.**
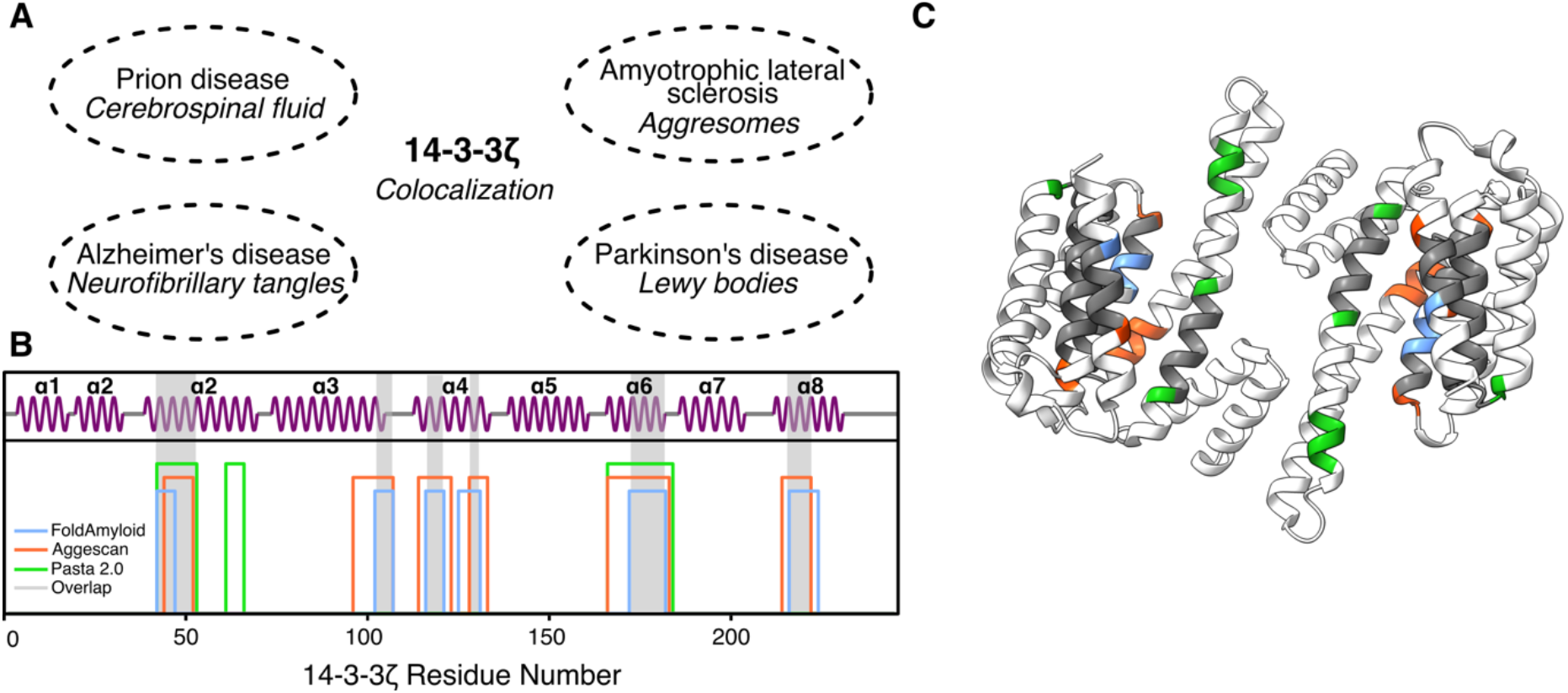
(**A**) 14-3-3ζ co-localization sites in various neurogenerative disorders: in cerebrospinal fluid in prion diseases^35,36^, in aggresomes during Amyotrophic lateral sclerosis^37^, in neurofibrillary tangles in Alzheimer’s disease ^14^ and in Lewy bodies during Parkinson’s disease^38^. (**B**) Predicted aggregation-prone regions by FoldAmyloid (Blue), AGGRESCAN (red) and Pasta 2.0 (green). An overlap with at least two predictors is shown with a grey bar. The secondary structure was depicted using Biotite^39^.(**C**) The predicted aggregation-prone sites are located in α-helices of 14-3-3ζ (PDB id: 5NAS). The protein structure was visualized with ChimeraX^40^

### 14-3-3ζ aggregates bind amyloid-specific dyes

For an initial aggregation control assay, the incubated 14-3-3ζ solution was combined with ThT and the sample fluorescence spectra were scanned as described in the Materials and Methods section. The resulting signal was quite low, which prompted the need for a higher sample concentration prior to the dye assays. The 14-3-3ζ aggregates were pelleted and resuspended into a 10 times lower volume before being used in the assays. Furthermore, in order to account for the possibility of non-amyloid fluorescence enhancement of ThT, three additional dye molecules were used, which included ANS (fluorescence increases in hydrophobic environments)^41^, Congo red (absorbance spectrum changes upon binding to amyloid fibrils)^42^ and a commercial dye – Amytracker 630, which recently has been shown to have very strong photophysical properties when binding to amyloid fibrils^43^.

When comparing the intensity of ThT in PBS or with the initial 14-3-3ζ solution, there were virtually no differences between the spectra (Fig. 2A). Over the seven days of incubation, we measured the 14-3-3ζ solution and observed a significant increase in the fluorescence emission intensity with a maximum at 485 nm (Fig. 2A, B). The increase in sample light scattering also confirmed the formation of larger structures (Fig. 2C). In the case of ANS, the aggregated 14-3-3ζ solution had the highest signal value (Fig. 2D), however, the initial 14-3-3ζ solution also displayed a considerable level of fluorescence. Similar results were obtained when using Amytracker630 (Fig. 2E), suggesting that 14-3-3ζ was capable of incorporating these molecules, with most likely them binding in the hydrophobic patch within the binding groove^4^. When CR was combined with the aforementioned solutions, only the incubated 14-3-3ζ sample displayed a significant change in the absorbance spectra (Fig. 2F), with the appearance of a shoulder at 540 nm. Such a shift is usually associated with the interaction between CR and amyloid fibrils, as is the increase in total absorbance in the 450 – 600 nm range^42^.

**Figure 2.**
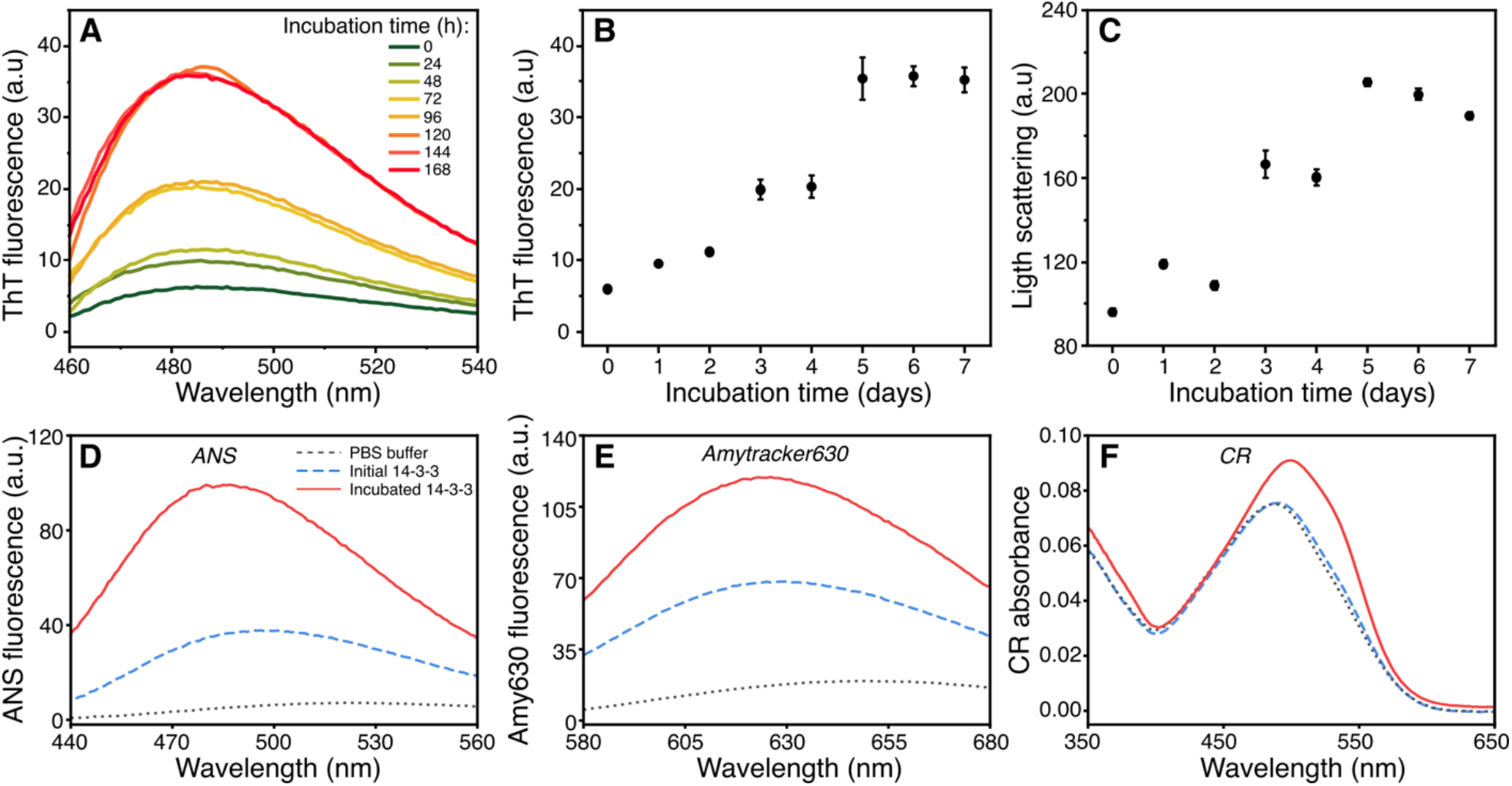
(**A**) The ThT fluorescence spectra of incubated 14-3-3ζ solution at different time points. The ThT fluorescence emission (**B**) and light scattering intensity (**C**) over a seven-day period. The ANS (**E**), Amytracker630 (**E**) fluorescence and CR (**F**) absorbance spectra of PBS buffer, initial 14-3-3ζ and incubated 14-3-3ζ (7 days) solution.

**Figure 3.**
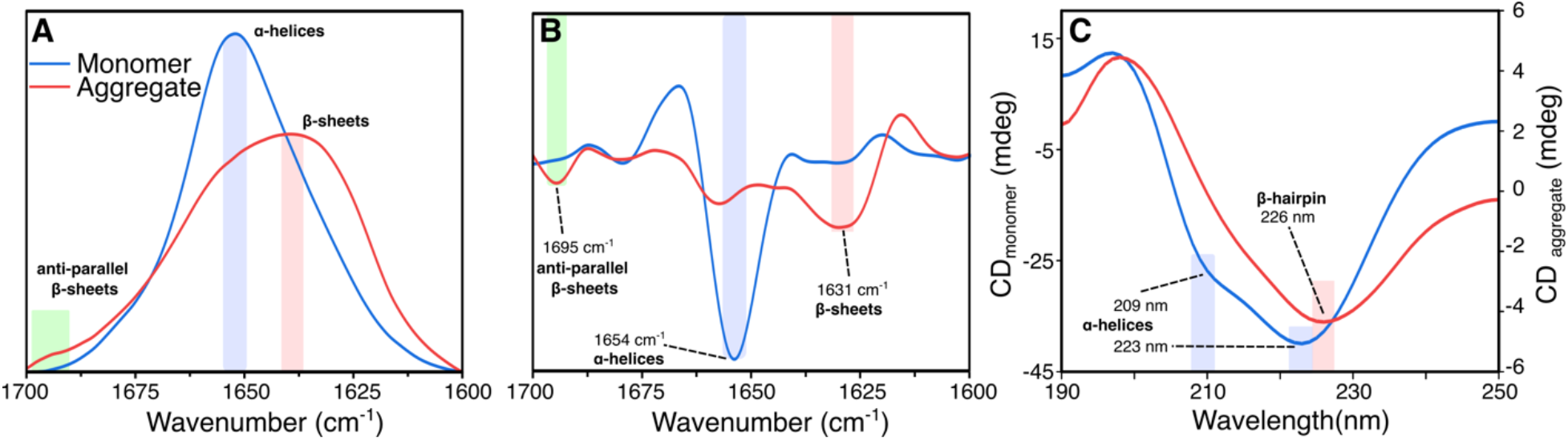
FTIR absorption (**A**), second derivative (**B**) and CD (**C**) spectra of monomeric and aggregate solution of 14-3-3ζ.

### Formation of β-sheets in 14-3-3ζ aggregate

In the case of monomeric 14-3-3ζ, the FTIR spectrum main maximum position was at 1651 cm^-1^ (associated with the presence of α-helical secondary structure^44^). When 14-3-3ζ was in its aggregated state, the FTIR spectrum main maximum position shifted towards 1638 cm^-1^ and we observed the appearance of an additional minor band at 1695 cm^-1^ (possibly anti-parallel β-sheets). The second derivative of the monomeric 14-3-3ζ FTIR spectrum had a main minimum at 1654 cm^-1^. Oppositely, the main minimum of the aggregated 14-3-3ζ FTIR spectrum second derivative was at 1631 cm^-1^ (associated with β-sheets) and two other clear minima at 1657 cm^-1^ (α-helices or turns) and 1695 cm^-1^ (anti-parallel β-sheets).

As complementary to FTIR data, we recorded CD spectra of 14-3-3ζ monomers and aggregates. The monomer spectrum had two minimum peaks at 209 nm and 223 nm, which is typical for α-helical proteins. On the other hand, the incubated sample spectra had only one unusual red-shifted minimum at 226 nm, which previously has been assigned to β-hairpins in other proteins and peptides^46,47^. This also matches with the observed FTIR band at 1695 cm^-1^, due to the fact that the anti-parallel β-sheet structure is a component of β-hairpins^48^.

### 14-3-3ζ aggregates resemble amyloid fibrils

As a final confirmation that 14-3-3ζ assembled amyloid fibrils, we used AFM imaging to observe the structure and shape of formed aggregates. The initial AFM image showed short amyloid fibrils evenly distributed over the surface of the mica (Fig. 4A). Upon closer inspection, we observed straight or slightly curved fibrils (Fig. 4B). The fibril height distribution was spread between 1-3 nm, with the mean being 1.6 ± 0.4 nm. The small shape of the fibrils resembled previously detected worm-like fibrils of the pro-inflammatory S100A9 protein^49^ and protofibrils of α-synuclein^50^.

**Figure 4.**
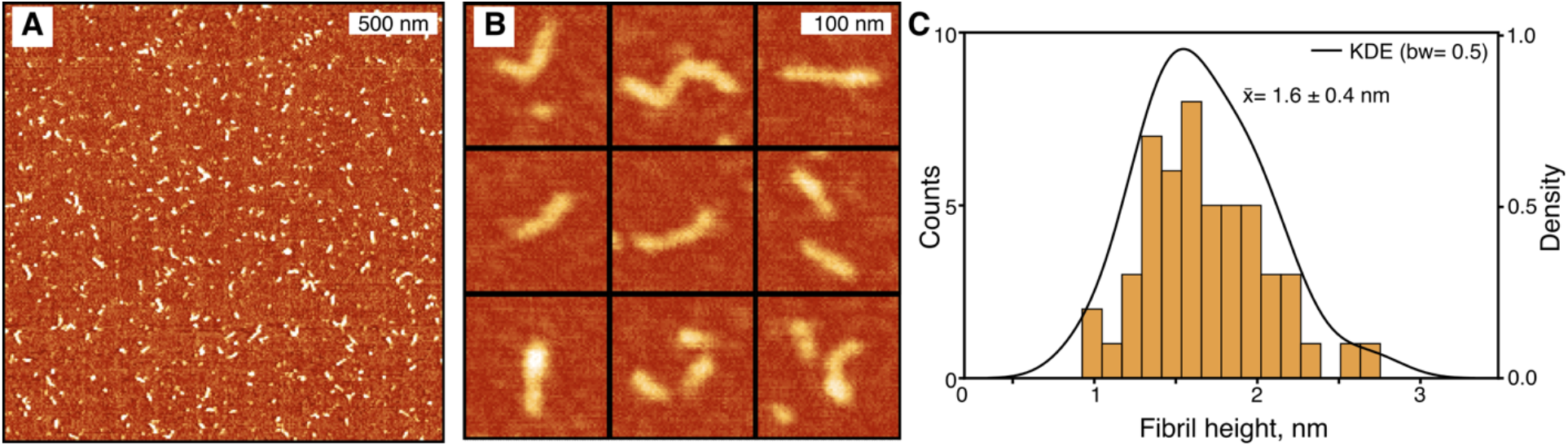
(A) AFM image of formed 14-3-3ζ amyloid fibrils (scale bar 500 nm). (B). Close-up view of fibrils showing curvy morphology. (C). Height distribution of fibrils. The black line is the kernel density estimation (KDE) function fit.

## Conclusions

In this study, for the first time, we observed the formation of amyloid fibrils by 14-3-3ζ. The small shape morphology and relatively weak fluorescence of bound amyloidogenic dyes indicate that 14-3-3ζ fibrils are not easily detectable, but behave similarly to amyloids. Since the detection of 14-3-3 proteins is often associated with the onset of neurodegenerative diseases^25,35,51^, our study suggests that this might not only be due to their neuroprotection^52^ and chaperone^53^ roles but also due to the formation of 14-3-3 fibrils. Moreover, with recent studies indicating 14-3-3 involvement in liquid-liquid phase separation, it is becoming evident that 14-3-3 might have an even larger role in neurodegenerative disorders than initially thought^54,55^. As it is now, the revelation of 14-3-3ζ amyloid properties opens up new possibilities for further investigations into other isoforms of 14-3-3 and their relationship to other protein aggregation pathways.

## Acknowledgements

The authors thank Björn Marcus Burmann and Jorge Eduardo Azevedo for gifting 6xHis-SUMO-14-3-3ζ and pET28a-HsSENP1 (Addgene plasmid #71465) plasmids accordingly.

## Funding

### Author Contribution

**Darius Šulskis**: Conceptualization, Methodology, Writing - Original Draft, Visualization. **Mantas Žiaunys**: Methodology, Writing -Review & Editing. **Andrius Sakalauskas**: Methodology, Writing -Review & Editing. **Vytautas Smirnovas**: Supervision, Writing -Review & Editing.

## Data and materials availability

The kinetic and FTIR, CD data used for analysis have been tabulated and are available on Mendeley Data: 10.17632/564277pjyx.1. All other relevant data are available from the corresponding author upon reasonable request.

