## Supplemental Information for "Formation of amyloid fibrils by the regulatory 14-3-3ζ protein"

### for

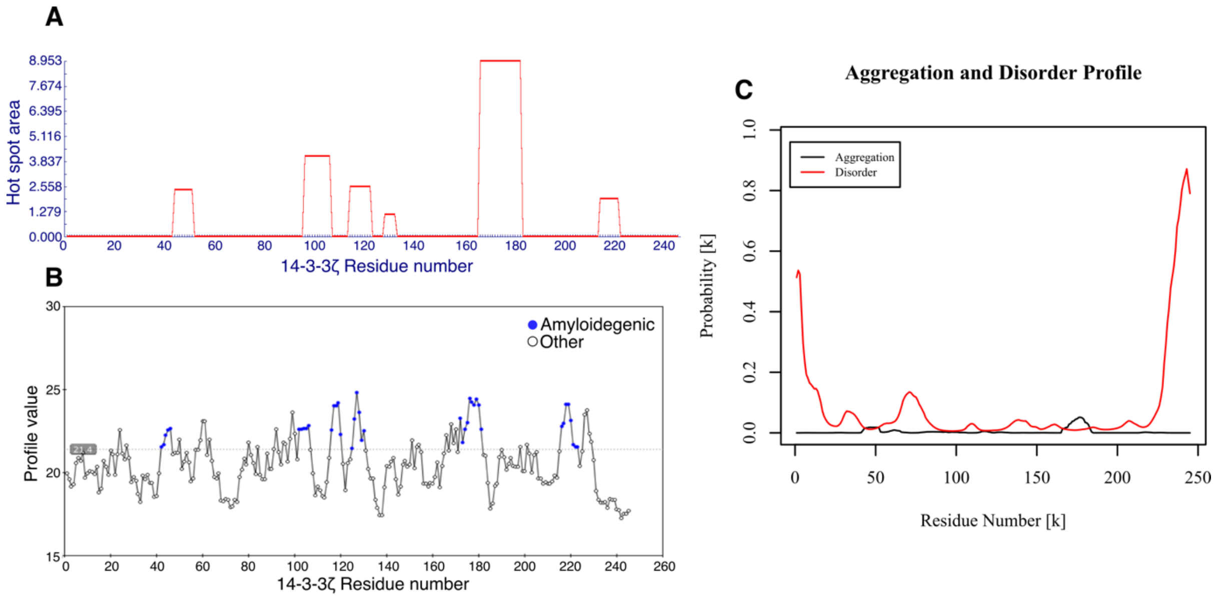


Figure S1.
Prediction of 14-3-3ζ aggregation-prone regions by AGGRESCAN (**A**), FoldAmyloid (**B**) and PASTA 2.0 (**C**)

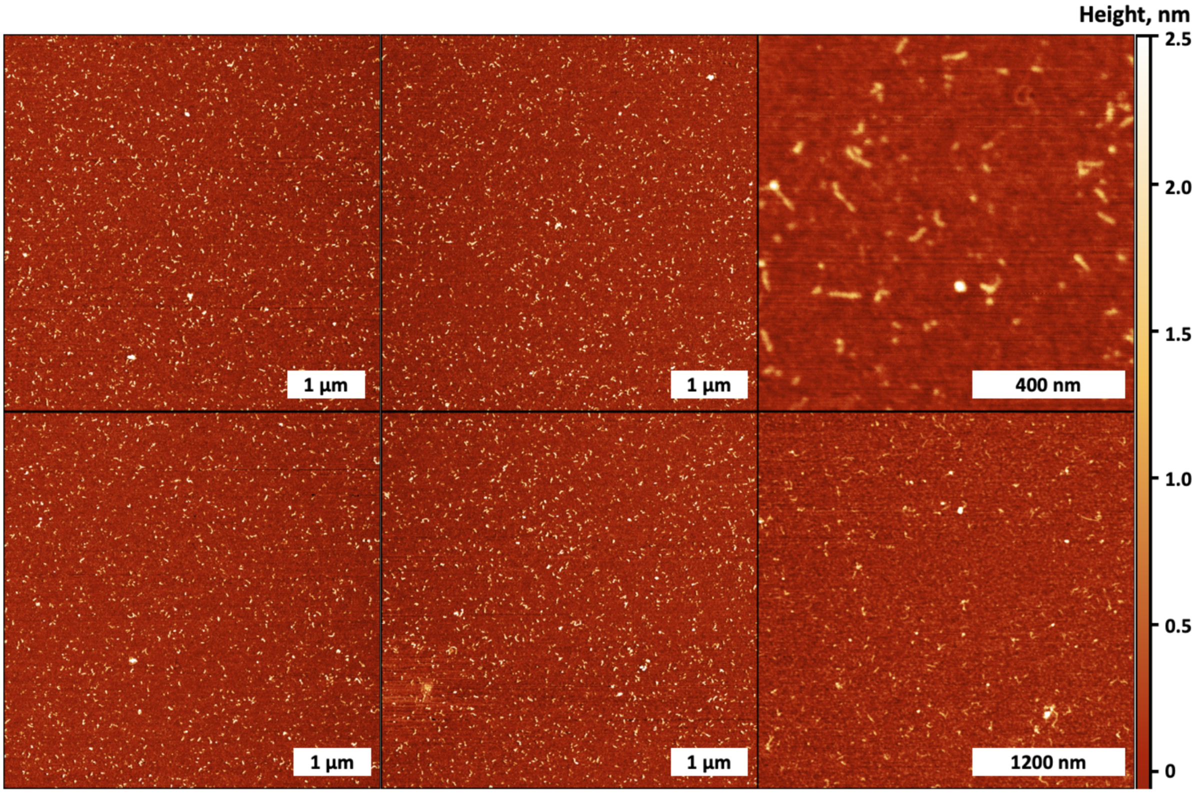


Figure S2.
AFM images of 14-3-3ζ aggregates.

| Plasmid | Primer | Sequence |
|---|---|---|
| pDS17<br>(6xHis-SUMO-14-3-3ζ ) | DS17_BamHI_Rev | 5’GTGGGATCCTTAATTTTCCCCTCCTTCTCC 3' |
| pIB01<br>(Senp1 catalytic domain 415-644 a.a.) | Senp1_del_frw<br>Senp1_del_rev | 5’ CCGGAAATTCGCCGCTGCTGTGATGATGATGATG 3’<br>5’CAGCAGCGGCGAATTTCCGGAAATCACCGAAGAAATG 3’ |

Supplementary Table 1: Plasmids and respective primers used in this study.
